# An internal sensor detects dietary amino acids and promotes food consumption in *Drosophila*

**DOI:** 10.1101/204453

**Authors:** Zhe Yang, Rui Huang, Xin Fu, Gaohang Wang, Wei Qi, Wei Shen, Liming Wang

**Affiliations:** Life Sciences Institute, Zhejiang University, Hangzhou, Zhejiang, China; Innovation Center for Cell Signaling Network, Zhejiang University, Hangzhou, Zhejiang, China; Institute of Neuroscience, Shanghai Institutes for Biological Sciences, Chinese Academy of Sciences, Shanghai, China; University of Chinese Academy of Sciences, Beijing, 100049, China; School of Life Science and Technology, ShanghaiTech University, Shanghai, 201210, China

**Keywords:** feeding, DH44, CRH, *Drosophila*

## Abstract

Adequate protein intake is crucial for animals. Despite the recent progress in understanding protein hunger and satiety in the fruit fly *Drosophila melanogaster*, how fruit flies assess prospective dietary protein sources and ensure protein consumption remains elusive. We show here that three specific amino acids, L-glutamate (L-Glu), L-alanine (L-Ala), and L-aspartate (L-Asp), rapidly promote food consumption in fruit flies when present in food. The effect of dietary amino acids to promote food consumption is independent of mating experience and internal nutritional status. Genetic analysis identifies six brain neurons expressing diuretic hormone 44 (DH44) as a sensor of dietary amino acids. DH44^+^ neurons can be directly activated by these three amino acids, and are both necessary and sufficient for dietary amino acids to promote food consumption. By conducting single cell RNAseq analysis, we also identify an amino acid transporter, CG13248, which is highly expressed in DH44^+^ neurons and is required for dietary amino acids to promote food consumption. Therefore, these data suggest that dietary amino acids may enter DH44^+^ neurons via CG13248 and modulate their activity and hence food consumption. Taken together, these data identify an internal amino acid sensor in the fly brain that evaluate food sources post-ingestively and facilitates adequate protein intake.

## INTRODUCTION

Proteins are the most abundant macromolecules in living organisms with a vast array of biological functions. Adequate and balanced protein consumption is therefore vital for the survival, reproduction, and well-being of animals. In fruit flies *Drosophila melanogaster*, several layers of regulatory mechanism are involved in the regulation of protein intake. Prolonged protein deprivation leads to feeding preference towards protein-rich diet and increased protein consumption (Liu et al., 2017; Ribeiro and Dickson, 2010; Toshima and Tanimura, 2012; Vargas et al., 2010). The detection of protein hunger and the induction of subsequent protein feeding involve a small group of dopaminergic neurons in the fly brain (Liu et al., 2017). Conversely, shortly after feeding on protein-rich diet, the insulin-producing cells (IPCs) in the fly brain are activated and exert robust suppressive effect on protein feeding (Liu et al., 2015; Wu et al., 2005). The activation of IPCs after protein intake is directly mediated by circulating L-Leucine (L-Leu) via a leucine transporter minidiscs (MND) and glutamate dehydrogenase (GDH) (Manière et al., 2016), and indirectly mediated by a fat body derived satiety hormone named female-specific independent of transformer (FIT) (Sun et al., 2017). In addition, flies rapidly detect and reject food sources lacking one or more essential amino acids (essential amino acid deficiency, EAAD), which is regulated by a different set of dopaminergic neurons (Bjordal et al., 2014). Collectively, these neural mechanisms ensure fruit flies to assess their internal amino acid adequacy, and to ensure sufficient and balanced intake of amino acids from desirable food sources.

It remains controversial, however, how fruit flies assess the presence of amino acids in potential food sources and modulate food consumption accordingly. In mammals, dietary amino acids elicit umami taste via the T1R1/T1R3 taste receptor located on the oral taste buds, which is believed to play a fundamental role in facilitating the evaluation and consumption of potential protein sources (Nelson et al., 2002). But fruit flies lack the homologs of mammalian umami taste receptor (Liman et al., 2014). A recent report has shown that specific taste neurons expressing Ir76b, an ionotropic chemosensory receptor, are sensitive to various amino acids to different extents (Ganguly et al., 2017). These taste neurons are located in female tarsi and may be involved in detecting amino acids in food sources. But it remains unclear whether Ir76b and Ir76b^+^ taste neurons play a direct role in initiating feeding behavior and ensure adequate protein consumption. Notably, eliminating *Ir76b* gene or silencing Ir76b^+^ neurons do not completely eliminate preference towards protein-rich diet, suggesting the presence of additional amino acid sensing mechanism(s) (Ganguly et al., 2017).

In this present study, we examined the effect of dietary amino acids to modulate food intake in fruit flies. We found that dietary amino acids significantly promoted food intake independent of flies’ mating experience and internal nutritional status. We also found that among all 20 natural amino acids, only three of them, including L-Glu, L-Ala, and L-Asp, but not their unnatural D-enantiomers, enhanced food consumption. In the fly brain, these three amino acids directly activated a small group of neurons expressing DH44, the homolog of mammalian corticotropin-releasing hormone (CRH). These DH44^+^ neurons are both necessary and sufficient for dietary amino acids and dietary yeast to promote food consumption.

We further investigated the cellular mechanism underlying the activation of DH44^+^ neurons by specific dietary amino acids. We identified CG13248, an amino acid transporter, highly expressed in DH44^+^ neurons and required for dietary amino acids to promote feeding. These results suggest that dietary amino acids directly enter DH44^+^ neurons and modulate their activity and hence food consumption. Taken together, we identify an internal sensor in the fly brain that rapidly detects specific dietary amino acids and promote protein feeding.

## RESULTS

### Dietary amino acids rapidly promote food consumption despite the lack of rapid taste responses

We first asked whether dietary amino acids could modulate food consumption. Since amino acids alone did not elicit food consumption (data not shown), we examined the effect of dietary amino acids on food consumption by adding varying concentrations of amino acids (from 5 mM to 200 mM) to 400 mM sucrose. By using the MAFE (<u>Ma</u>nual <u>Fe</u>eding) assay that measured the volume of ingested food by individual flies during the course of a single meal (Figure 1a) (Qi et al., 2015), we found that the presence of amino acids significantly increased food consumption of virgin female flies fed *ad libitum* (Figure 1b) without altering flies’ proboscis extension reflex (PER) responses (Figure 1c). The increase in food consumption was likely a result of increased feeding duration in the presence of dietary amino acids (Figure 1d). To understand whether amino acids alone could promote feeding or they merely enhanced feeding towards sucrose, we examined water consumption in dry-starved flies. These thirsty flies consumed significantly more water in the presence of amino acids (Figure 1e), indicating that dietary amino acids alone are capable of promoting feeding behavior. Yeast is the major protein source for fruit flies in their natural habitats and in laboratory conditions (Grandison et al., 2009; Spieth, 1974). We also found that adding 5% yeast in 400 mM sucrose significantly promoted food consumption (Figure 1f). Thus, dietary amino acids rapidly promote food consumption despite the lack of peripheral taste responses.

**Figure 1.**
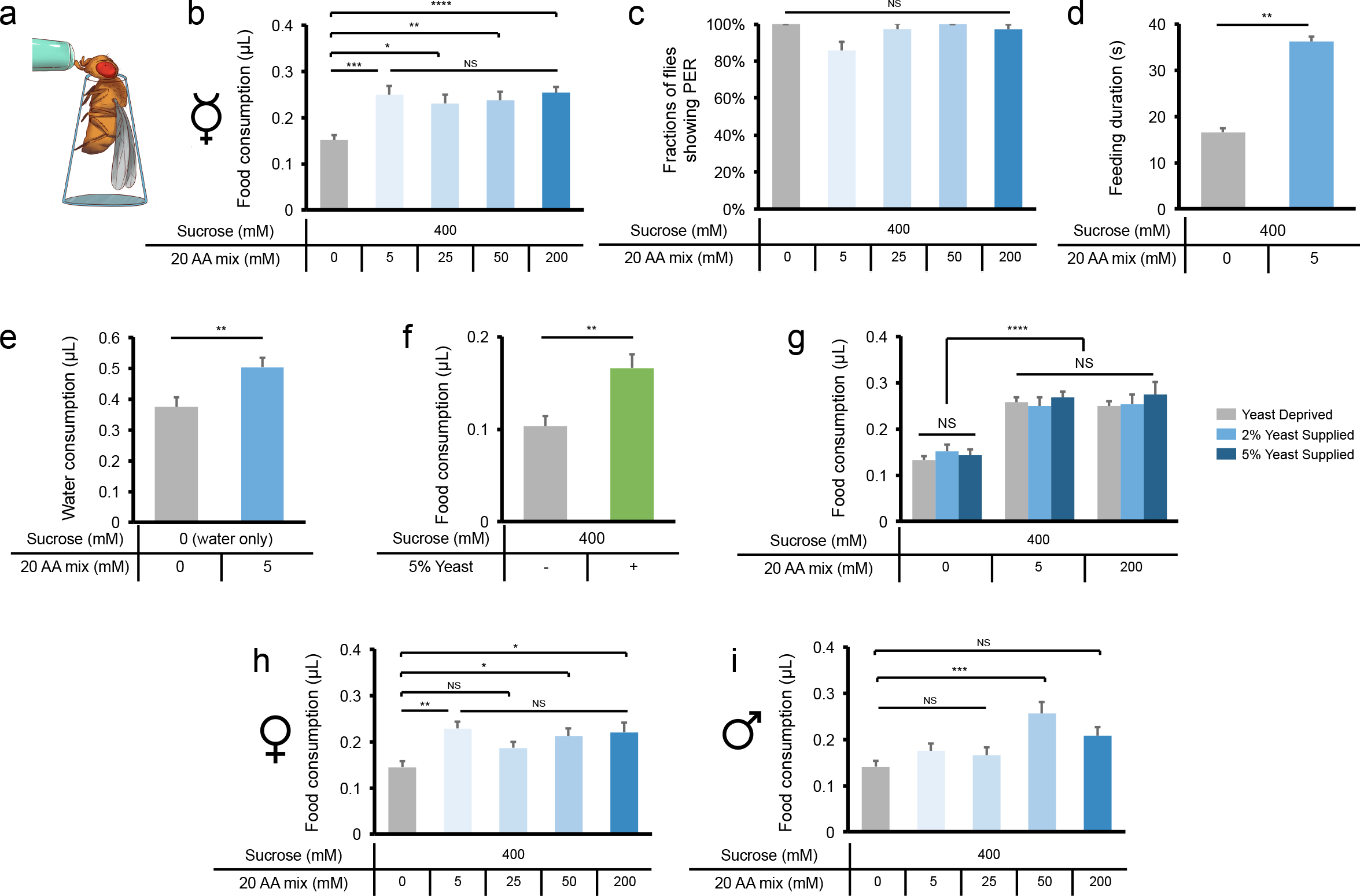
Dietary amino acids promote food consumption. (**a**) Schematic illustration of the MAFE assay. (**b**) Volume of 400 mM sucrose (grey) or 400 mM sucrose plus different concentrations of amino acid mixture (blue) consumed in a meal by *Canton-S* virgin females fed *ad libitum* with 5% sucrose plus 2% yeast (n=21-59). The composition of the amino acid mixture is shown in Table S1 (Grandison et al., 2009). (**c**) Fractions of *Canton-S* virgin females showing PER responses to sucrose alone (grey) or sucrose plus different concentrations of amino acid mixture (blue) (n=38-40). (**d**) The feeding duration of *Canton-S* virgin flies when fed with sucrose alone (grey) or sucrose plus amino acid mixture (blue) (n=20). (**e**) Volume of water consumption in the presence or absence of 5 mM amino acid mixture by dry-starved *Canton-S* virgin females (n=28-32). (**f**) Volume of food consumed by *Canton-S* virgin flies, in the presence or absence of 5% yeast extract (n=17-18). (**g**) Volume of food consumed by *Canton-S* virgin flies that were raised in the absence of yeast (“Yeast Deprived”), in the presence of 2% yeast, or 5% yeast (n=18-47). (**h-i**) Volume of food consumed by mated *Canton-S* females (**h**) or males (**i**) fed *ad libitum* with 5% sucrose plus 2% yeast (n=18-24). Data are shown as means (± SEM). NS, P > 0.05; *P < 0.05; **P < 0.01; ***P < 0.001; ****P < 0.0001.

Feeding preference towards protein-rich diet of fruit flies is dependent on the internal nutritional state (Ganguly et al., 2017; Liu et al., 2017; Ribeiro and Dickson, 2010; Sun et al., 2017; Vargas et al., 2010). We therefore tested whether complete protein deprivation also modulated the effect on food consumption by dietary amino acids. Yeast-deprived flies and those raised in the presence of 2% and 5% yeast exhibited comparable food consumption in the absence or presence of amino acids (Figure 1g), suggesting that similar to mammalian umami sensing, dietary amino acids enhance feeding independent of flies’ internal nutritional status. Notably, previous reports have also shown that female flies’ mating experience significantly enhanced their preference towards yeast (Ribeiro and Dickson, 2010; Vargas et al., 2010). Nevertheless, we found that dietary amino acids enhanced feeding to similar extents in both virgin (Figure 1b) and mated females (Figure 1h). Dietary amino acids also enhanced feeding of male flies, although to a lesser extent (Figure 1i).

### Three specific amino acids, L-Glu, L-Ala, and L-Asp, promote food consumption

Mammalian umami sensing neurons detect most natural L-amino acids but with differing sensitivities (Nelson et al., 2002). In fruit flies, Ir76b^+^ neurons located on fly tarsi are also responsive to most if not all natural L-amino acids and enhance yeast preference (Ganguly et al., 2017). Since dietary amino acids promoted food consumption independent of peripheral taste responses, we thus sought to identify the amino acids that mediated such effect in fruit flies. We found that out of twenty natural L-amino acids, only three of them, L-Glu, L-Ala, and L-Asp, significantly enhanced food consumption (Figure 2a).

**Figure 2.**
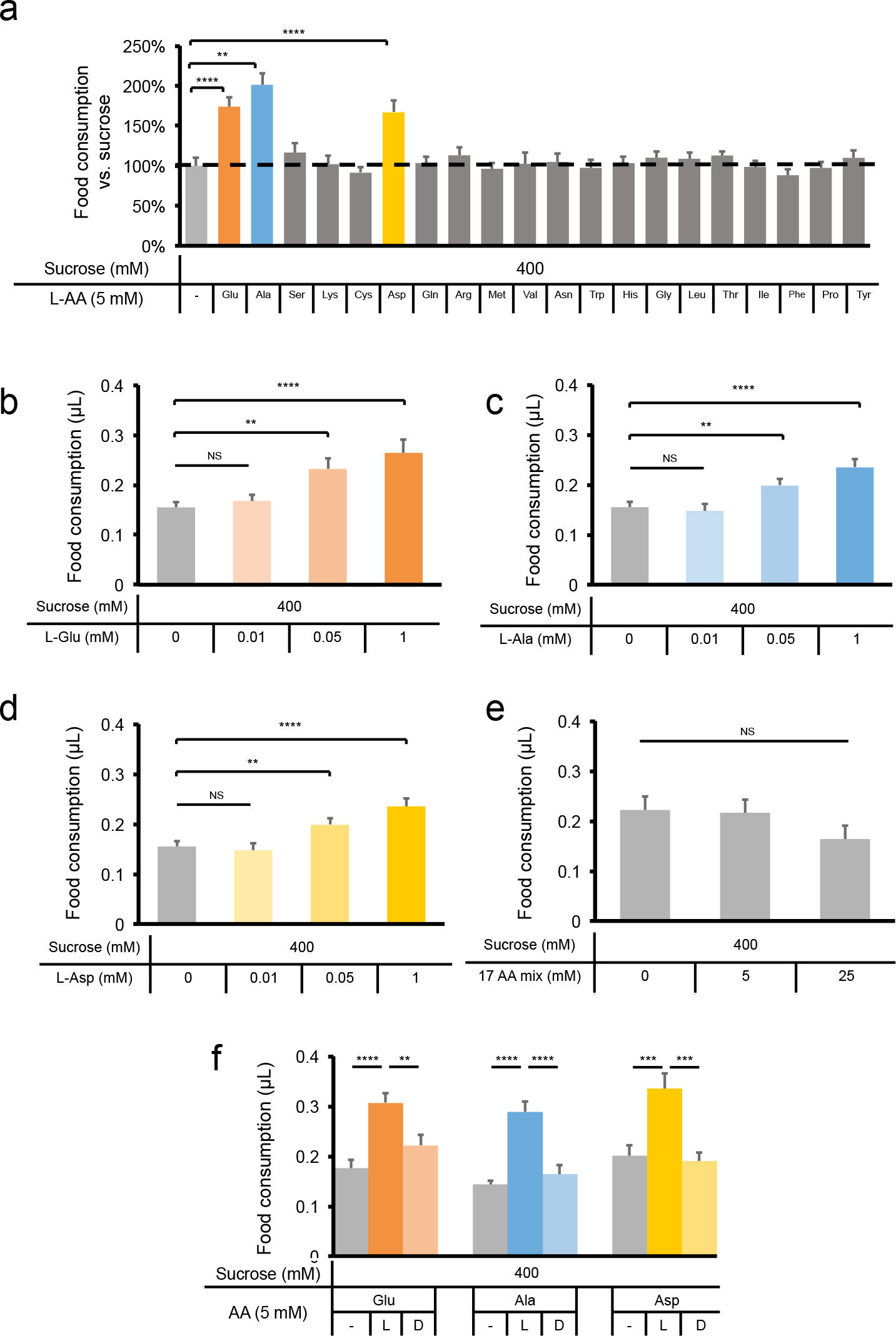
Three specific amino acids, L-Glu, L-Ala, and L-Asp, promote food consumption. (**a**) Volume of 400 mM sucrose or 400 mM sucrose plus 5 mM amino acid consumed by *Canton-S* virgin females (n=20-26) (except for tyrosine, for which we used its saturated concentration 2.7 mM). (**b-d**) Volume of 400 mM sucrose plus different concentrations of L-Glu (**b**), L-Ala (**c**), and L-Asp (**d**) consumed by *Canton-S* virgin females (n=15-37). (**e**) Volume of 400 mM sucrose plus different concentrations of the other 17 amino acids consumed by *Canton-S* virgin females (n=13-15). (f) Volume of 400 mM sucrose or 400 mM sucrose plus 5 mM L-or D-amino acid consumed by *Canton-S* virgin females (n=20-26). Data are shown as means (± SEM). NS, P > 0.05; **P < 0.01; ***P < 0.001; ****P < 0.0001.

These three L-amino acids promoted food consumption in a dose-dependent manner, starting at concentrations equal to or higher than 0.05 mM (Figure 2b-d). In contrast, the mixture of the other 17 amino acids did not promote food consumption even at much higher concentrations (Figure 2e, 5 and 25 mM). Similar to mammalian umami taste, only the L-amino acids, but not their unnatural D-enantiomers, enhanced food consumption in fruit flies (Figure 2f).

### DH44 signaling is required for dietary amino acids to promote food consumption

Dietary amino acids enhanced food consumption without altering flies’s PER responses. Therefore, a post-ingestive amino acid sensor in the central nervous system might mediate the feeding promoting effect of dietary amino acids. To identify the putative amino acid sensor, we screened a collection of candidate neuropeptide receptors by using the MAFE assay (Figure 3a). Among the receptors we screened, knocking down DH44 Receptor 1 (DH44R1) and leucokinin receptor (LKR) in the nervous system by using a pan-neuronal GAL4 driver, abolished the effect of dietary amino acids to enhance food consumption. LKR signaling has been shown to regulate different aspects of feeding behavior, including the meal size and feeding frequency (Al-Anzi et al., 2010) and sleep regulation after protein intake (Murphy et al., 2016). Therefore in this present study we focused on the function of DH44 signaling in amino acid sensing and food consumption.

**Figure 3.**
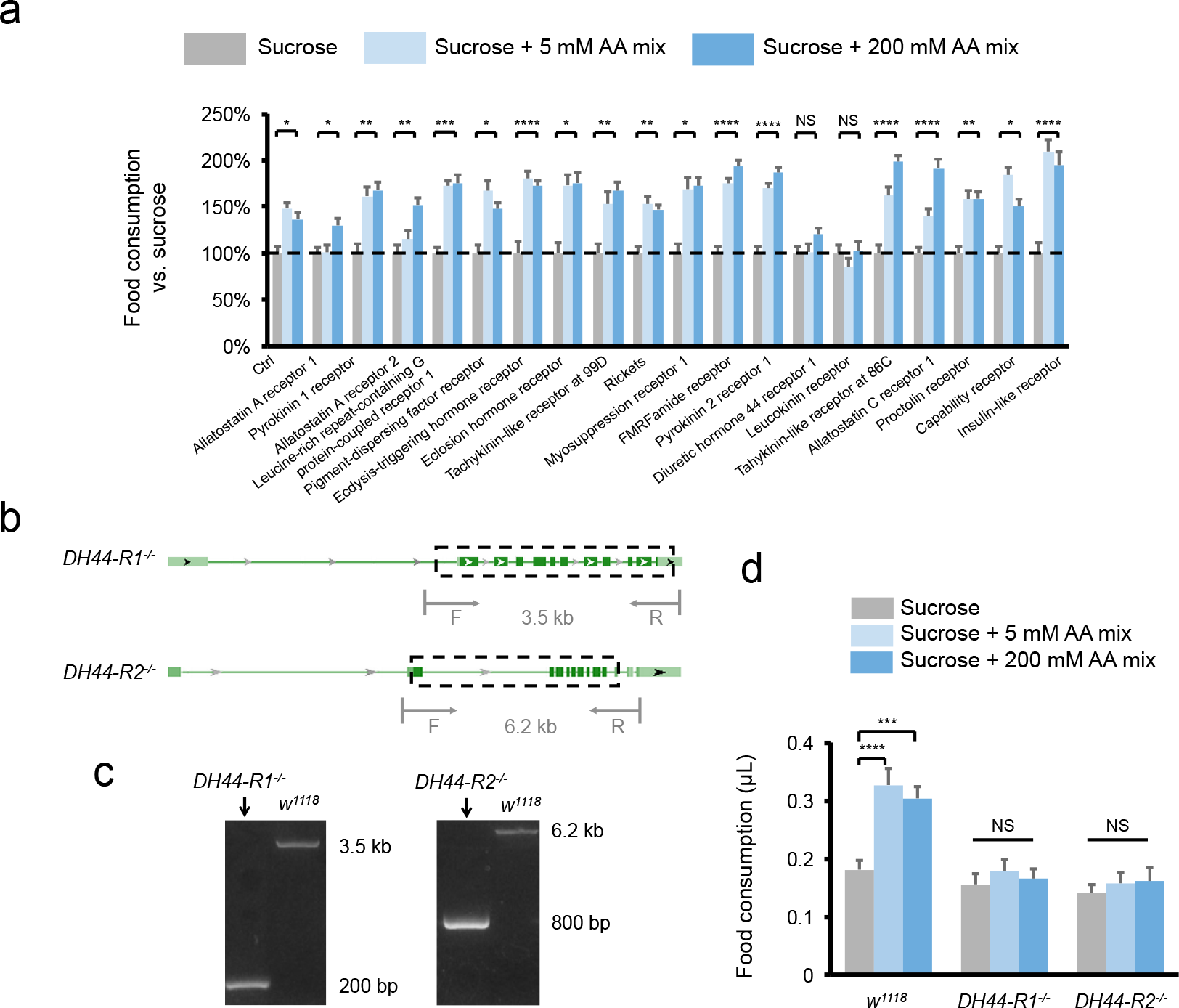
DH44 signaling mediates the effect of dietary amino acids to promote feeding. (**a**) Changes in food consumption to 400 mM sucrose plus different concentrations of amino acid mixture consumed by flies with pan-neuronal RNAi knockdown of indicated neuropeptide receptors (n=19-35). Food consumption to 400 mM sucrose alone is used as the baseline (grey, dotted line). (**b**) Genomic organization of *DH44R1* and *DH44R2* genes. Dotted boxes outline the deleted DNA fragments by CRISPR/cas9. The positions of PCR primers for genotyping are also illustrated. (**c**) Validation of *DH44R1*^-/-^ and *DH44R2*^-/-^ mutants by genomic PCR. (**d**) Volume of 400 mM sucrose (grey) or 400 mM sucrose plus different concentrations of amino acid mixture (blue) consumed by indicated genotypes (n=19-23). Data are shown as means (± SEM). NS, P > 0.05; **P < 0.01; ***P < 0.001; ****P < 0.0001.

DH44 is the fly homolog of mammalian CRH (Lovejoy and Jahan, 2006). Like mammals, fly DH44 has two receptors, DH44R1 and DH44R2 (Figure 3b) (Johnson et al., 2003). DH44R1 is expressed in the nervous system and is involved in feeding control, whereas DH44R2 is expressed in gut enteroendocrine cells and regulates gut motility and excretion (Dus et al., 2015). We generated genetic mutants for both DH44 receptors, DH44R1 and DH44R2, via CRISPR/cas9 mediated gene editing (Figure 3b-c). Both *DH44R1*^-/-^ and *DH44R2*^-/-^ mutants exhibited completely abolished responses to dietary amino acids (Figure 3d). These results suggest that DH44 signaling is required for dietary amino acids to promote food consumption.

### DH44^+^ neurons are directly activated by dietary amino acids

DH44 is specifically expressed in six neurosecretory cells in the pars intercerebralis (PI) region of the fly brain (Figure 4a-b) (Dus et al., 2015). It has been shown that DH44^+^ neurons could be directly activated by nutritive sugars including D-glucose and D-fructose, suggesting that DH44^+^ neurons are a post-ingestive nutrient sensor in the fly brain (Dus et al., 2015). We thus asked whether DH44^+^ neurons were also responsive to dietary amino acids. To this aim, we performed calcium imaging on *ex vivo* brain preparations (Figure 4c-d). Similar preparation has reliably recorded calcium transients in these neurons as described previously (Dus et al., 2015). Consistent with the behavioral responses (Figure 2), perfusion of L-Glu, L-Ala, and L-Asp activated DH44^+^ neurons in a dose-dependent manner (Figure 4e-g). The D-enantiomers of these three amino acids elicited significantly lower calcium responses than the L-enantiomers at the same concentration (Figure 4e-g). In addition, several L-amino acids that were unable to promote food consumption, including L-glutamine (L-Gln), L-methionine (L-Met), and L-asparagine (L-Asn), induced much smaller calcium transients (Figure 4h).

**Figure 4.**
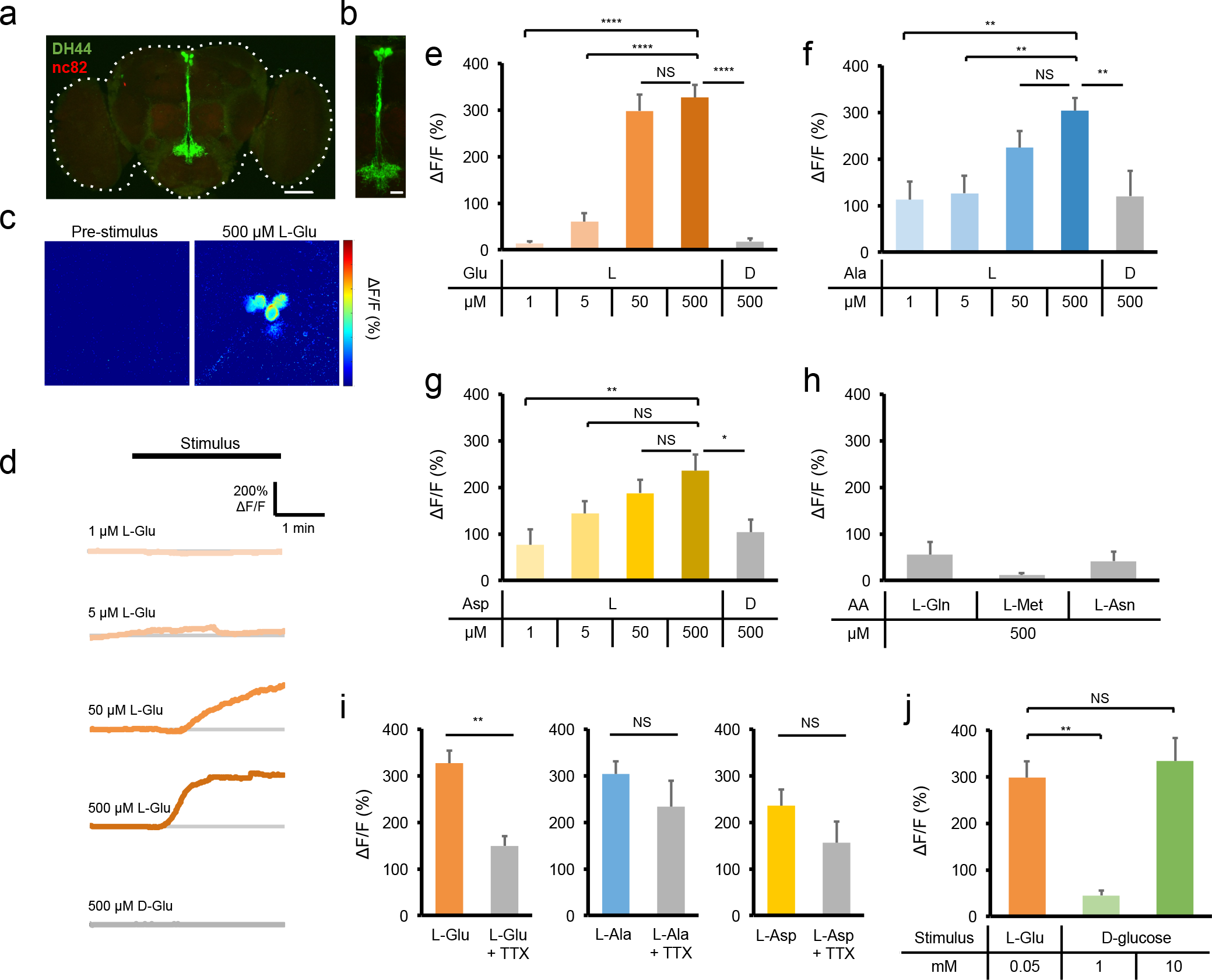
DH44^+^ neurons are directly activated by dietary amino acids. (**a**) The expression of membrane bound mCFD8GFP in DH44^+^ neurons. The scale bar represents 20 μm. The dotted line outlines the fly brain. (**b**) An enlarged image of the PI region seen in (**a**). The scale bar represents 10 jim. Green: mCD8GFP in DH44^+^ neurons. Red: nc82. (**c**) The calcium responses of DH44^+^ neurons to 500 μM L-Glu (right). (**d**) Representative traces of the calcium responses of DH44^+^ neurons to different concentrations of L-Glu and D-Glu. (**e-h**) Quantification of the calcium responses of DH44^+^ neurons to different concentrations of indicated amino acids (n=9-40). (**i**) Quantifications of the calcium responses of DH44^+^ neurons to 500 μM of indicated amino acids in the absence or presence of 1 μM TTX (n=10-22). (**j**) Quantifications of the calcium responses of DH44^+^ neurons to L-Glu and D-glucose (n=9-15). Data are shown as means (± SEM). NS, P > 0.05; *P<0.05; **P < 0.01; ****P < 0.0001.

To test whether DH44^+^ neurons were directly sensitive to amino acids, we used tetrodotoxin (TTX), a blocker of voltage-gated Na^+^ channels, to eliminate synaptic inputs of DH44^+^ neurons. We found that TTX did not affect calcium transients elicited by L-Ala and L-Asp, and a small effect in reducing calcium responses to L-Glu (Figure 4i). Thus, the results of TTX blockage suggest that DH44^+^ neurons are a direct sensor to dietary amino acids.

It is worth noting that DH44^+^ neurons are much more sensitive to dietary amino acids than nutritive sugars. In DH44^+^ neurons, 0.05 mM L-Glu elicited much higher calcium transients than 1 mM D-glucose and comparable calcium responses to 10 mM D-glucose (Figure 4j).

### DH44^+^ neurons mediate the effect of dietary amino acids to promote food consumption

We next asked whether DH44^+^ neurons mediated the effect of dietary amino acids to promote feeding. Genetic silencing of DH44^+^ neurons by ectopic expression of Kir2.1 (Nitabach et al., 2002), an inwardly rectifying potassium channel, did not affect sucrose consumption (Figure 5a). In the presence of L-Glu, L-Ala, and L-Asp, however, silencing DH44^+^ neurons significantly reduced food consumption (Figure 5b). Therefore DH44^+^ neurons are required for the increase in food consumption by dietary amino acids but not dietary sugars (Figure 5 c).

**Figure 5.**
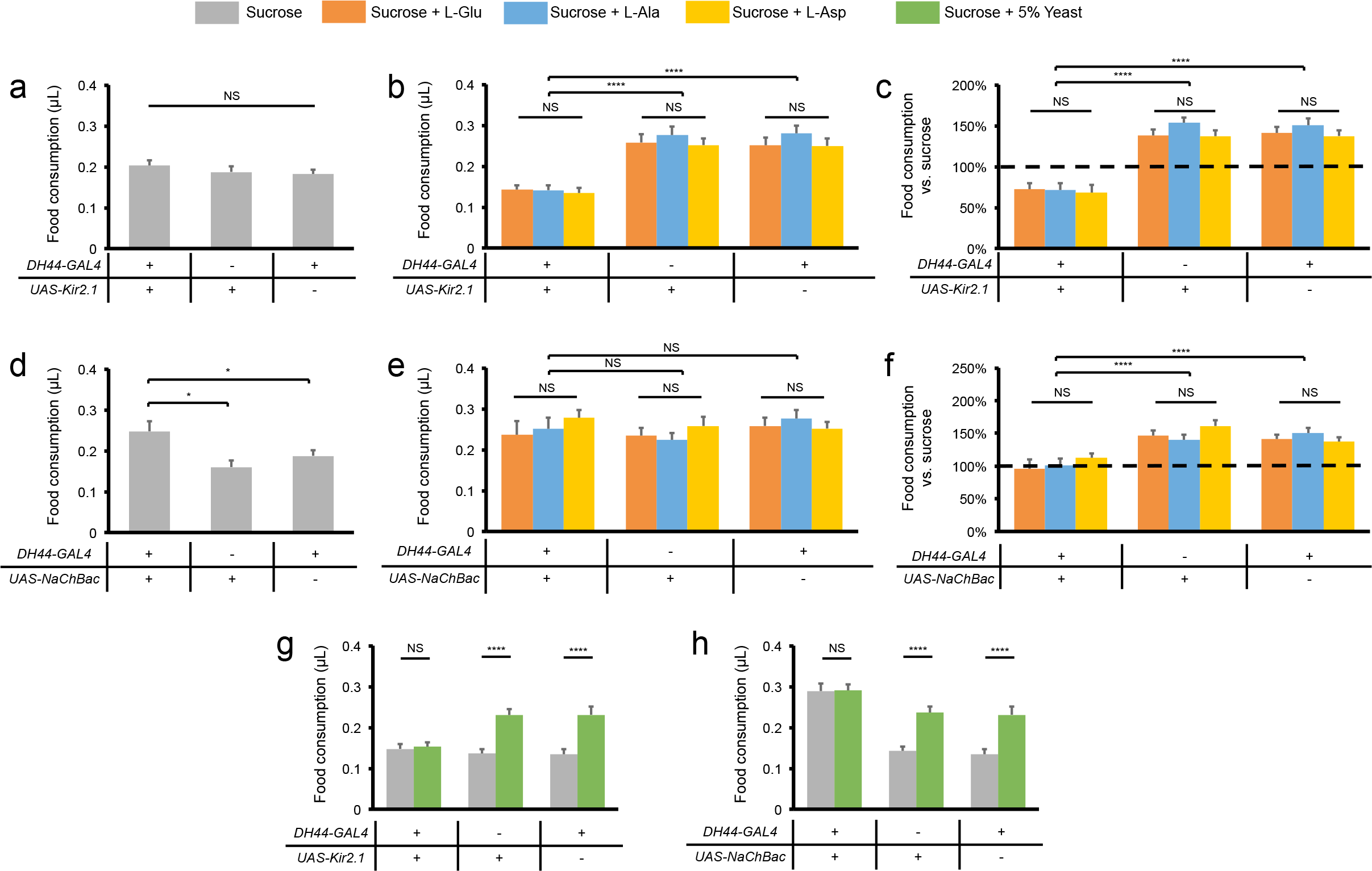
DH44^+^ neurons mediate the effect of dietary amino acids and yeast to promote feeding. (**a, d**) Volume of 400 mM sucrose consumed by indicated genotypes (n=21-65). (**b, e**) Volume of 400 mM sucrose plus 5 mM of indicated amino acid consumed by indicated genotypes (n=20-38). (**c, f**) Changes in food consumption by the addition of 5 mM of indicated amino acid compared to 400 mM sucrose alone (dotted line) (n=20-38). (**g, h**) Volume of 400 mM sucrose consumed in the presence or absence of 5% yeast extract by indicated genotypes (n=23-36). Data are shown as means (± SEM). NS, P > 0.05; ****P < 0.0001.

In contrast, artificial activation of DH44^+^ neurons by NaChBac (Nitabach et al., 2005), a bacterial sodium channel, significantly enhanced the consumption of sucrose (Figure 5d). DH44^+^ neuronal activation, however, did not further increase the consumption of dietary amino acids (Figure 5e). Therefore, activation of DH44^+^ neurons mimicked the effect of dietary amino acids to promote food consumption (Figure 5f).

Taken together, our results suggest that DH44^+^ neurons mediate the effect of dietary dietary amino acids to promote food consumption, which may help to ensure adequate protein intake of flies. Consistently, DH44^+^ neurons also mediated the effect of yeast, the major protein source of fruit flies, to promote food consumption (Figure 5g-h).

### Dietary amino acids enter DH44^+^ neurons via an amino acid transporter CG13248

As previously suggested, nutritive sugars may need to enter DH44^+^ neurons and modulate their activity (Dus et al., 2015). It is possible that dietary amino acids may also need to enter DH44^+^ neurons to modulate food consumption. We thus sought to identify amino acid transporter(s) that mediate the entry of dietary amino acids into DH44^+^ neurons. We conducted single-cell RNAseq experiments of individual DH44^+^ neurons following a previously described protocol (Yu et al., 2016). Among ~50 candidate amino acid transporters (Limmer et al., 2014), 17 of them showed expression in more than half of the DH44^+^ neurons we examined (Figure 6a). We were able to obtain RNAi or genetic mutant reagents for 9 out of these 17 genes (Figure 6a, red). Among them, eliminating CG13248 and CG4991 blocked the effect of dietary amino acids to promote food consumption (Figure 6b).

**Figure 6.**
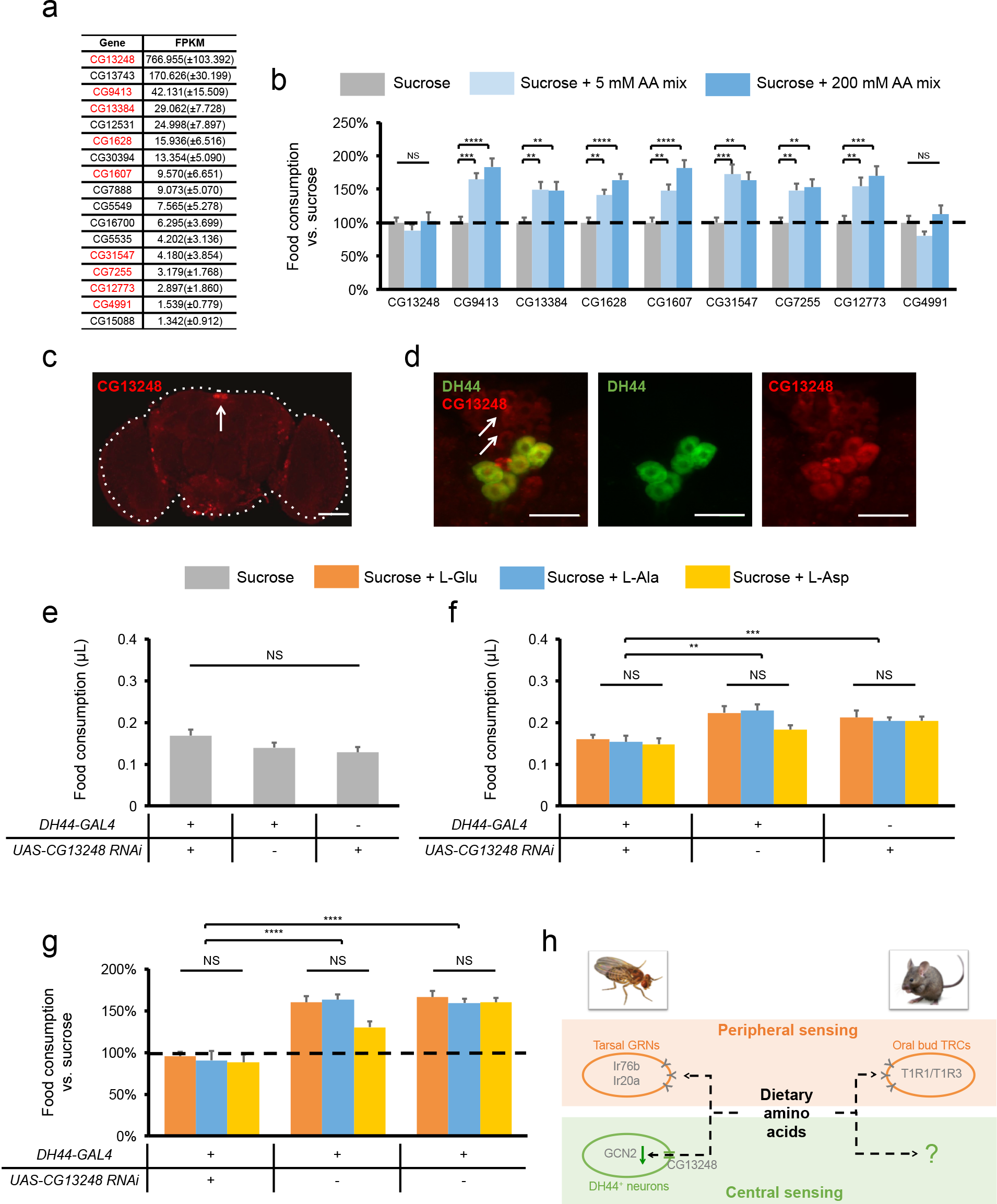
Dietary amino acids may enter DH44^+^ neurons via an amino acid transporter CG13248. (**a**) The list of putative amino acid transporter genes expressed in DH44^+^ neurons. All genes that are expressed in more than 50% of individual DH44^+^ neurons (FPKM≥1) are listed here. Genes in red indicate the presence of genetic reagents to evaluate their behavioral function. (**b**) Changes in food consumption by the addition of amino acid mixture compared to 400 mM sucrose alone (dotted line) when indicated gene are eliminated either by genetic mutation (CG31547) or by panneuronal RNAi knock-down (all other genes) (n=20-23). See Materials and Methods for details. (**c**) Expression of CG13248 in the fly brain. The scale bar represents 20 μm. Red: CG13248. The dotted line outlines the fly brain. The arrow indicates the PI region. (**d**) Co-localization of CG13248 and DH44 in the PI region of the fly brain. The scale bar represents 10 μm. Green: mCD8GFP in DH44^+^ neurons. Red: CG13248. The arrows indicate neurons with CG13248 expression but not DH44 expression. (**e**) Volume of 400 mM sucrose consumed by indicated genotypes (n=21-23). (**f**) Volume of 400 mM sucrose plus 5 mM of indicated amino acid consumed by indicated genotypes (n=20-26). (**g**) Changes in food consumption by the addition of 5 mM of indicated amino acid compared to 400 mM sucrose alone (dotted line) (n=20-26). (**h**) Detection of dietary amino acids in rodents vs. in fruit flies. In rodents, a specific group of taste-receptor cells (TRCs) located on the taste buds are responsive to most L-amino acids via T1R1/T1R3 taste receptor and facilitate food intake. In flies, besides the peripheral amino acid sensing mechanism (tarsal gustatory sensory neurons (GRNs) expressing both Ir76b and Ir20 receptors), the present study uncovers a central censing mechanism that DH44^+^ neurons detect L-Glu, L-Ala, and L-Asp by suppressing GCN2 signaling and promote food consumption. Data are shown as means (± SEM). NS, P > 0.05; **P<0.01; ***P<0.001; ****P < 0.0001.

CG13248 is a cationic amino acid transporter (Park et al., 2011). Antibody against CG13248 revealed its expression in the fly brain, particularly in the PI region (Figure 6c, arrow). In that region, CG13248 was expressed in ~10 neurons per fly brain, including all six DH44^+^ neurons (Figure 6d). The co-localization of CG13248 and DH44 further suggests that CG13248 may mediate amino acid sensing in DH44^+^ neurons. Consistently, knocking down CG13248 expression specifically in DH44^+^ neurons did not interfere with sucrose consumption (Figure 6e), but blocked the effect of L-Glu, L-Ala, and L-Asp to promote feeding (Figure 6f-g).

Notably, SLC7A4, the human homolog of CG13248, was insufficient to confer amino acid transporter activity when heterologously expressed (Wolf et al., 2002). It is therefore plausible that additional proteins, including other amino acid transporters, may be required for the entry of L-Glu, L-Ala, and L-Asp into DH44^+^ neurons. Evidently, we found that besides CG13248, CG4991 was also required for L-Glu, L-Ala, and L-Asp to promote food consumption (Figure 6b and Figure S1). Whether and how these two transporters work together for amino acid entry remains to be further elucidated.

## DISCUSSION

In this present study, we have used a previously development feeding assay, named the MAFE assay, to study the regulation of food consumption in fruit flies (Qi et al., 2015). In the MAFE assay, individual flies are immobilized when liquid food is administrated directly to their proboscis. As we previously reported, the regulation of food seeking, feeding initiation, and food consumption are independent. Therefore, a unique advantage of the MAFE assay is that it measures food consumption without the interference from the process to search for food and the initiation of food intake. A potential caveat of the MAFE assay is that flies’ feeding behavior may has a “forced” component since food is present directly to their mouthpart. But as we have previously characterized, the meal size in the MAFE assay is comparable to that of free-moving flies in the CAFÉ assay (Ja et al., 2007; Qi et al., 2015).

By using the MAFE assay, we have demonstrated that dietary amino acids significantly promote food consumption. It is worth noting that unlike flies’ feeding preference towards yeast (Ribeiro and Dickson, 2010; Vargas et al., 2010), the effect of dietary amino acids to promote feeding is independent of flies’ mating experience and internal nutritional state. Therefore, the sensing of dietary amino acids may be specifically involved in the assessment of potential food sources but not flies’ internal amino acid adequacy.

Among all twenty natural amino acids, only three of them, L-Glu, L-Ala, and L-Asp, but not their D-enantiomers nor the other seventeen L-amino acids, exhibit such effect. Notably, L-Glu, L-Ala, and L-Asp are among the most abundant amino acids in yeast, the major protein source of fruit flies. Therefore we speculate that the putative sensor of dietary amino acids in flies is tuned to these three specific amino acids, which provides a reliable assessment of protein-rich food sources for fruit flies and facilitates adequate protein intake. Evidently, the detection thresholds of these three amino acids (0.05 mM) are several magnitudes lower than their concentrations in yeast (~30-50 mM) (Grandison et al., 2009) and in synthetic medium optimized for flies’ lifespan and fecundity (~3-5 mM) (Piper et al., 2014).

We have identified an internal sensor in the fly brain that detects these three specific dietary amino acids and promotes food consumption in response. Behavioral, genetic, and functional imaging studies collectively reveal that DH44^+^ neurons in the fly brain can be directly activated by low concentrations of L-Glu, L-Ala, and L-Asp, but not by their D-enantiomers or all other L-amino acids. The activation of DH44^+^ neurons then sustains the feeding activity and increases food consumption within the time course of a single meal. These properties of DH44^+^ neurons may facilitate the search, evaluation, and consumption of desirable protein-rich food sources and ensure protein homeostasis in fruit flies.

Previous work has shown that DH44^+^ neurons detect nutritive sugars in food including D-glucose and D-fructose (Dus et al., 2015). It is possible that DH44^+^ neurons function as a universal post-ingestive sensor that evaluates various types of nutrients in the gastrointestinal (GI) tract. Notably, DH44+ neurons are significantly more sensitive to amino acids than to sugars. Given that in flies’ food sources, the concentrations of sugars (~50 mM) are also significantly higher than amino acids ( ~3-5 mM) (Piper et al., 2014), this unique property of DH44^+^ neurons may help to ensure the detection of both classes of nutrients in food sources and agile regulation of food intake behavior.

At the cellular level, we have identified CG13248, an amino acid transporter, highly expressed in DH44^+^ neurons and required for their function in promoting food consumption. It is therefore likely that dietary amino acids can directly enter these neurons and modulate their neuronal activity. Notably, the axonal projections of DH44^+^ brain neurons innervate extensively with the GI tract of flies, including the esophagus and the gut (Dus et al., 2015). It is therefore plausible that dietary nutrients may penetrate the blood brain barrier and enter DH44^+^ neurons along the GI tract (Limmer et al., 2014). Detailed anatomical characterization of DH44^+^ neurons will be required to further elucidate this important problem.

As strict heterotrophs, rapid detection of amino acids in potential food sources is critical for adequate and balanced intake of protein diet. In mammals, amino acid sensing occurs peripherally, by T1R1/T1R3 taste receptor on the oral taste buds (Efeyan et al., 2015; Nelson et al., 2002). In fruit flies, two distinct amino acid sensing mechanisms exist. Peripherally, Ir76b^+^ taste neurons located on the legs are responsive to most L-amino acids and mediate yeast preference in protein-starved flies (Ganguly et al., 2017). And centrally, DH44^+^ neurons in the fly brain detect post-ingestive L-Glu, L-Ala, and L-Asp and enhance food consumption (Figure 6h). It is of interest to understand the potential crosstalk between these two mechanisms in securing adequate protein intake. Notably, protein-starved flies exhibit reduced yet considerable yeast preference in the absence of *Ir76b* gene or Ir76b^+^ neurons (Ganguly et al., 2017). One possible explanation is that in the absence of peripheral amino acid sensing, DH44^+^ neurons can still sense the presence of dietary amino acids post-ingestively and regulate food intake.

The identification of an internal amino acid sensor in fruit flies also raises the question of whether similar post-ingestive amino acid sensing mechanism exists in mammals (Figure 6h). Besides the peripheral umami sensing, mammalian brain can detect the presence of free amino acids in the circulation system. For example, specific hypothalamic neurons are responsive to circulating L-Leu via TOR signaling and suppress food intake (Cota et al., 2006; Mori et al., 2009). In addition, EAAD sensing in rodents is also taste-independent and involves the anterior piriform cortex (Hao et al., 2005; Leib and Knight, 2015; Maurin et al., 2005). However, unlike DH44^+^ neurons in the fly brain, these known internal amino acid sensing mechanisms in mammals play a suppressive role in food intake. It will be of interest to explore the presence of taste-independent mechanism that detects dietary amino acids and promotes food intake in mammals.

## MATERIALS AND METHODS

### Flies

Flies were kept in vials containing standard fly medium made of yeast, corn, and agar at 25□ and 60% humidity and on a 12-h light-12-h dark cycle. If not otherwise indicated, virgin female flies were collected shortly after eclosion and kept on medium made of 5% sucrose and 2% yeast before subjected to behavioral experiments. Dry-starved flies were kept in vials with no food or water supply for 18-24 hours before behavioral experiments.

All *UAS-RNAi* lines used in the neuropeptide receptgor screen (#25858, #25935, #25936, #27275, #27280,#27506,#27509,#27513,#27529, #27539, #28780, #28781, #28783, #29414, #31884, #31958, #35251, #36303, #38346, #38347), *UAS-TOR^ted^* (#7013), and *elav-GAL4* (#25750) were obtained from the Bloomington *Drosophila* Stock Center at Indiana University. For the amino acid transporter screen, the genetic mutant for CG31547 was obtained from the Bloomington *Drosophila* Stock Center at Indiana University (#59219). For other candidate amino acid transporter genes, *UAS-RNAi* lines were obtained from the Tsinghua Fly Center (#3116, #02064.N, #4215, #02284.N, #04347.N, #04422.N, #04964.N, #01189.N2). *DH44-GAL4* was from Yi Rao (Peking University). *UAS-GCN2 RNAi* was from Pierre Léopold (Université de Nice, INSERM) and *UAS-ATF4 RNAi* was from the Tsinghua Fly Center.

*DH44RV*^-/-^ and *DH44RZ*^-/-^ mutant flies were generated using the CRISPR/cas9 system. Briefly, a pair of sgRNA (*DH44R1*: GTCAATTGTTAGGGGATTCCCCGG, GTTGTAAAATACTTGAAGCAGTGG and GTTTGCTCCGCTTGGTACTTGG and GTTCATAGCATGGAGTTGGTTGG; *DH44R2*: GAAGTGCCAGAGTTCAGGAGTGG and GTTCATTCATAGTGTCCAGTGGG) were co-injected together with cas9 mRNA. Germline transmissions were confirmed by PCR amplication and Sanger sequencing (*DH44R1*: forward primer-TAAGCCGAGTTCGATGTG, reverse primer-CCATTTGCACATTGAGTTAC; *DH44R2*: forward primer-CACACTCGTGCCAACTAA, reverse primer-AAGGACGCAGACAGATAAC). ~3.3 kb in *DH44R1* gene locus (14381876 to 14385244 in Chromosome 2R) and ~5.4 kb and *DH44R2* gene locus (12480822 to 12486265 in Chromosome 2R) were removed including most of the exons. Both mutants were viable and showed no apparent morphological and behavioral deficits.

### Chemicals

Sucrose (S7903), agar (A1296), L-alanine (A7627), L-argine (A5131), L-asparagine (A0884), L-aspartate (A8949), L-cysteine (C1276), L-glutamate (G1251), L-glutamine (G3126), L-glycine (G7126), L-histidinie (H8000), L-isoleucine (I2752), L-leucine (L8912), L-methionine (M9625), L-phenylalanine (P2126), L-proline (P0380), L-serine (S4500), L-theronine (T8625), L-tryptophan (T0254), L-tyrosine (T3754), L-valine (V0500), L-lysine (L5626), D-alanine (A7377), D-aspartate (V900627), D-glutamate (G1001), and denatomium (D5765) were purchased from Sigma-Aldrich. Yeast extract (LP0021) was purchased from Thermo Fisher Scientific.

### Behavioral assays

PER was assayed as described previously (Qi et al., 2015). Briefly, individual flies were gently aspirated and immobilized in a 200 μL pipette tip. Flies were first sated with water and then subjected to different tastants, each tastant tested twice. Flies showing PER responses to at least one of the two trials were considered positive to the tastant.

The MAFE assay was performed as described previously (Qi et al., 2015). Individual flies were transferred and mobilized as the PER assay. They were then presented with liquid food filled in a graduated glass capillary (VWR, 53432-604) until they stopped responding to food stimuli for ten serial food stimuli. Food consumption was calculated based on the volume change before vs. after feeding.

### Calcium imaging

Adult fly brains were freshly dissected into adult hemolymph-like solution (AHL) (108 mM NaCl, 8.2 mM MgCl_2_, 4 mM NaHCO_3_, 1 mM NaH_2_PO_4_, 2 mM CaCl_2_, 5 mM KCl, 5 mM HEPES, 80 mM sucrose, pH = 7.3) and immobilized with fine pins on a Sylgard-based perfusion chamber. The brain was recorded in the first minute to establish a base line. Then the solutions were changed to (AHL + amino acids, pH adjusted back to 7.3) for more than 5 minutes. Solutions in the perfusion chamber were controlled by a valve commander (Scientific instruments). All imaging was performed on a LSM 710 Carl Zeiss confocal microscope. The scanning resolution was 512 × 512 pixels, the optical zoom was 2×. One frame is 0.97s. Image analyses were performed in ImageJ and plotted in Excel (Microsoft) or Matlab (MathWorks). The ratio changes were calculated using the following formula: ΔF/F = [F – F_0_] / [F – background], where F is the mean florescence of cell body, F_0_ is the base line (one minute interval before stimuli), and background is the mean florescence of an area without tissues in the same frame as F is measured. 1 μM TTX was applied to the bath to block neural transmission.

### Single-cell RNAseq

As described previously (Yu et al., 2016), individual DH44^+^ cells (visualized by GFP experession) were harvested from dissected fly brain under a fluorescence microscope with a glass micropipette pulled from thick-walled borosilicate capillaries (BF120-69-10, Sutter Instruments). Individual cells were immediately transferred to lysate buffer and underwent reverse transcription and cDNA amplification (SMARTer Ultra Low RNA Kit for Sequencing, Clonetech). The amplified cDNA were sonicated to ~250 bp fragments by the Covaris S2 system and then subjected to end-repair, dA-tail, adaptor ligation, and 12 cycles of PCR amplification using the library preparation kit (NEBNext Ultra II DNA Library Prep Kit, NEB). The cDNA libraries were sequenced by Illumina Hiseq 2500 platform. The sequenced raw data were first pre-processed to remove low-quality reads, adaptor sequences and amplification primer. Reads were mapped to *Drosophila* genome and mapped reads were selected for further analysis. FPKM (Fragments Per Kilobase Of Exon Per Million Fragments Mapped) was used to quantify gene expression.

### Immunohistochemistry

Fly brains were dissected in PBS on ice and transferred to 4% PFA for fixation for 55 min. Fixed brains were incubated with Dilution/Blocking Buffer (10% Calf Serum and 0.5 % Triton X-100 in PBS) for 1.5-2 hr at room temperature and incubated in Dilution Buffer with primary antibodies at 4°C for 24-36 hr. These brains were then washed with Washing Buffer (0.5% Triton X-100 in PBS) for 4 × 15 min at room temperature and subsequently incubated in secondary antibodies for 23-24 hr at 4°C. The brains were washed three times with Washing Buffer again before mounted in Fluoroshield (Sigma-Aldrich). Samples were imaged with Nikon 10×/0.45 and 40×/0.80w. Antibodies were used at the following dilutions: mouse anti-nc82 (1:200, DSHB), rabbit anti-GFP (1:500, Life Technologies), Guinea pig anti-CG13248 (1:1000, a gift from Paul Taghert), Alexa Fluor 546 goat anti mouse (1:300, Life Technologies), Alexa Fluor 488 goat anti rabbit (1:300, Life Technologies), Alexa Fluor 594 goat anti guinea pig (1:500, Abcam).

### Statistical analysis

Data presented in this study were verified for normal distribution by D’Agostino-Pearson omnibus test. Student’s t test (for pairwise comparisons) and one-way ANOVA (for comparisons among three or more groups) were used. If one-way ANOVA detected a significant difference among groups, a *post hoc* test with Bonferroni correction was performed for multiple comparisons. Two-way ANOVA (and *post hoc* test if applicable) was applied for comparisons with more than one variant.

## ACKNOWLEDGEMENTS

We thank all lab members for helpful discussions and technical assistance. We thank Pierre Léopold, Zhefeng Gong, Paul Taghert, Yi Rao, and the Bloomington *Drosophila* stock center at Indiana University for fly stocks and reagents. We thank Shen Xian Hui for insightful discussions throughout the course of this study and Yan Li, Xiaoke Chen, Liqun Luo, and Dengke Ma for helpful comments on the manuscript. Danping Chen and Ye Wu provide scientific and administrative support in the laboratory. This study was funded by the National Natural Science Foundation of China (No. 31522026 for L.W. and No. X-0402-14-002 for W.S.) and the Thousand Young Talents Plan (L.W. and Y.S.).

---

**Figure S1. CG4991 is also required for dietary amino acids to promote food consumption**

(**a**) Volume of 400 mM sucrose consumed by indicated genotypes (n=19-28). (**b**) Volume of 400 mM sucrose plus 5 mM of indicated amino acid consumed by indicated genotypes (n=18-30). (**c**) Changes in food consumption by the addition of 5 mM of indicated amino acid compared to 400 mM sucrose alone (dotted line) (n=18-30). Data are shown as means (± SEM). NS, P > 0.05; *P<0.05; **P<0.01; ***P<0.001; ****P<0.0001.

**Table S1. Composition of amino acid mixture**

Concentrations of each of the twenty L-amino acids in 5 mM amino acid mixture.

